# The contribution of alternative splicing probability to the coding expansion of the genome

**DOI:** 10.1101/048124

**Authors:** Fernando Carrillo Oesterreich, Hugo Bowne-Anderson, Jonathon Howard

## Abstract

Alternative splicing results in the inclusion or exclusion of exons in an RNA, thereby allowing a single gene to code for multiple RNA isoforms. Genes are often composed of many exons, allowing combinatorial choice to significantly expand the coding potential of the genome. How much coding potential is gained by alternative splicing and what is the main contributor: alternative-splicing-depth or exon-count? Here we develop a splice-site-centric quantification method, allowing us to characterize transcriptome-wide alternative splicing with a simple probabilistic model, enabling species-wide comparison. We use information theory to quantify the coding potential gain and show that an increase in alternative splicing probability contributes more to transcriptome expansion than exon-count. Our results suggest that dominant isoforms are co-expressed alongside many minor isoforms. We propose that this solves two problems simultaneously, that is, expression of functional isoforms and expansion of the transcriptome landscape potentially without a direct function, but available for evolution.

## Introduction

The central dogma of gene expression states that information flows from gene to protein via an RNA intermediate, pre-mRNA, which contains coding sequences (exons) interrupted by non-coding sequences (introns). Splicing joins exons and removes introns from the pre-mRNA, yielding a mature transcript (Wahl et al., 2009). A splicing event is classified as constitutive if two exons in a pre-mRNA are always spliced together. A splicing event is classified as alternative if a given exon can be spliced to distinct exons. A typical pre-mRNA contains several potential alternative exons, leading to the possibility that many distinct mature transcripts are generated from a single gene by a combinatorial choice of exons. This can lead to the expansion of the cell’s repertoire of RNA molecules, the transcriptome (Graveley, 2001). The majority of human genes are indeed reported to be alternatively spliced (Wang et al., 2008) and alternative splicing is responsible for transcriptome differences between tissues, during development and in disease (Boutz et al., 2007; Brown et al., 2014; Irimia and Blencowe, 2012; Kornblihtt et al., 2013; Wang et al., 2008). An important open question is the extent to which alternative splicing leads to expansion of the transcriptome. In this manuscript we address this question by deriving a mathematical definition of extent which is based on experimental data.

Alternative splicing leads to approximately 150,000 distinct transcripts generated from all human genes (Modrek and Lee, 2002), a ratio of about 7.5 transcripts per gene, assuming the current gene count (Ezkurdia et al., 2014). This expansion serves to increase the coding potential of the genome. One way to quantify alternative splicing is simply to use this ratio. However, it is not clear whether simply counting distinct transcripts (as illustrated in Figure 1A) is the best way to quantify alternative splicing. This is because a dominant isoform together with a very rare isoform is weighted equally to a pair of isoforms that are both expressed with the same probability. Yet from an information point of view it makes more sense if the latter pair is weighted more than the former. This example shows the limitation of simply counting isoforms as a method of quantifying alternative splicing. Another way to quantifying alternative splicing is to measure the frequency of inclusion of individual exons (Barbosa-Morais et al., 2012; Graveley, 2001; Merkin et al., 2012; Wang et al., 2008). A limitation of this method is that it assumes a specific type of alternative splicing in which exons are either included with their flanking exons or omitted. What is needed is a quantification of alternative splicing that that makes no assumptions about the type of splicing yet is able to take into account the number of transcripts and their frequency of use.

**Figure 1.**
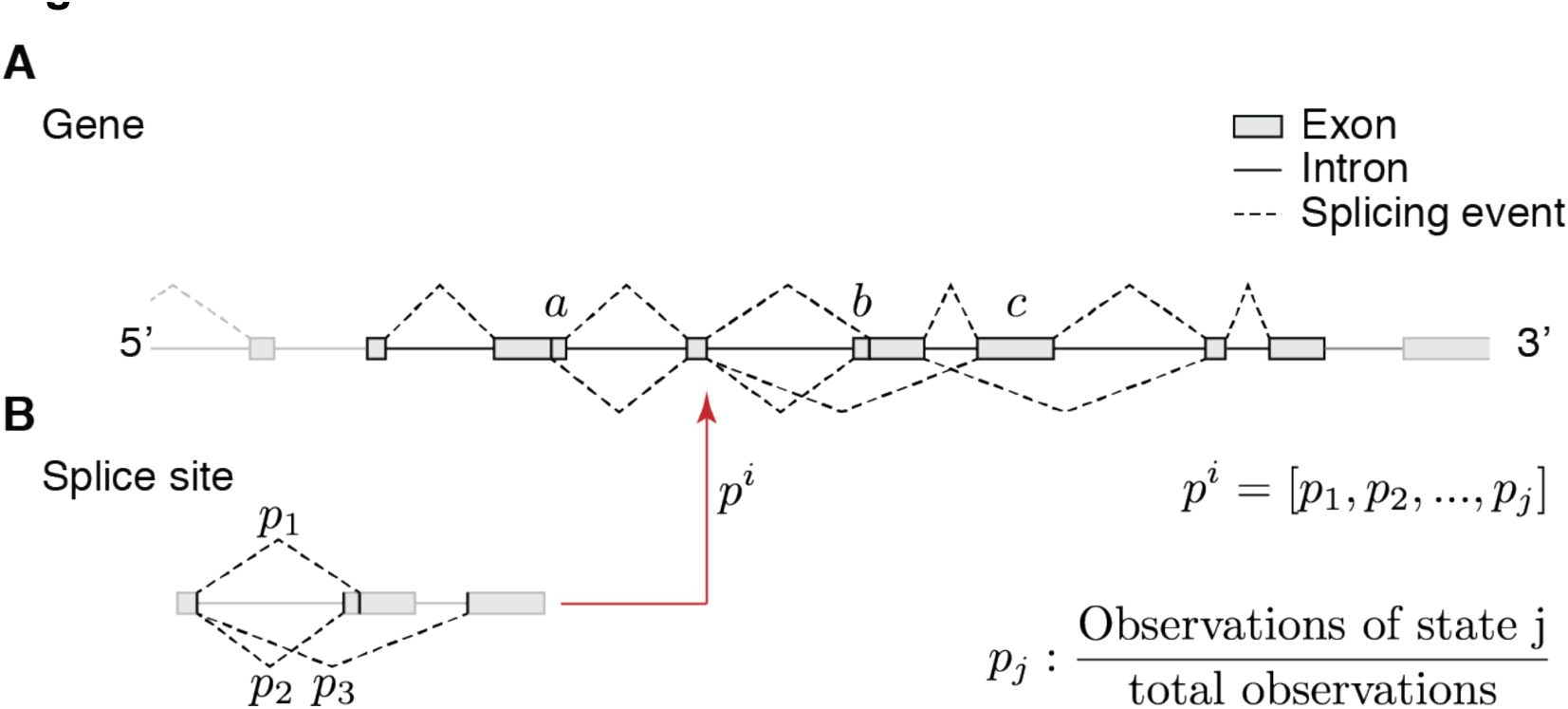
Gene and splice site centric views of alternative splicing. **(A)** Schematic representation of alternative splicing for a hypothetical gene. Exons (grey boxes), introns (solid lines) and splicing events (dashed lines) are indicated. The combinatorial nature of multiple splicing events can generate complex alternative splicing patterns for genes. Alternative splicing types represented here are alternative 3’ splice sites (a), alternative 5’ splice sites (b) and cassette exons (c). **(B)** A splice site centric approach significantly reduces complexity. One of the multiple splice sites of the gene shown in (A) is highlighted (left panel). This splice site is characterized by its accessible alternative splicing states and the number of observations for each splicing state, represented by the probability vector *p*^*i*^ (right panel). Transcriptome-wide splice site characterizations are derived from RNA-Seq experiments. Probability vector representation allows comparison of splice-sites independent of alternative splicing type.

To quantitate alternative splicing without making assumptions about the combinatorial complexity of splicing, we introduce a new splice-site centric measure of alternative splicing (Figure 1B and see Results section). This approach can be readily applied to transcriptome data for which the associated genome sequence is available and does not require that genome being annotated. We use this approach to analyze transcriptomes obtained by RNA-Seq from several different tissues, species and cells. By analyzing these data using our splice-site specific measure we show that (i) there is a simple model for the distribution of alternative splicing probabilities, (ii) that there are genomic features implicated in the regulation of alternative splicing that hold for all species analyzed and (iii) the probability of alternative splicing makes a larger contribution to transcriptome expansion than increasing exon count.

## Results

### Splice site centric view of alternative splicing

Alternatively spliced isoforms are the result of combinations of alternative splicing events (Figure 1A). Each splicing event joins two splice sites, defined as the boundaries between exon-intron and intron-exon sequences. We can therefore characterize splicing of a transcript by characterizing all its splice sites (Figure 1B). Each splice site has a number of splicing states (i.e. the partner splice sites that are used) and we characterize the splice site by the probabilities with which each state is chosen. In this scheme, alternative splicing is defined when a splice site has two or more splicing states.

We have used this splice site centric approach to analyze transcriptomes from large scale RNA-Seq studies. In these data, splicing events are represented by sequencing reads spanning joined exon-exon sequences, identifying splice-site pairs. We use the information contained in the sequencing reads to first deduce the set of all splice sites. For each splice site, we determined the number of splicing states (*N*_*i*_), counted the number of reads associated with each splicing state and divided each read by the total number of reads for that site. This yields a splicing probability distribution (a multinomial distribution) for each splice site (Figure 1B).

To identify characteristics of alternative splicing that may generalize across transcriptomes, we analyzed 23 publicly available RNA-Seq experiments obtained from 11 different species (see Supplementary Table 1). These transcriptomes were derived from widely evolutionarily distributed organisms, including algae, nematode worms, flies, mice and humans. Within species, data were obtained from single cell types or from complex mixtures of tissues. The data therefore span a wide range of expected transcriptome complexity. We quantified splice site choice individually for each splice site present in each transcriptome, represented by probability distributions.

To investigate how our splice site centric approach relates to the exon-centric approach discussed in the Introduction, we plotted the number of splice sites detected by our algorithm against the number of annotated exons (Figure 2A). The number of splice sites ranged from approximately hundred thousand (*D. melanogaster*) to several hundred thousand (*Mus musculus, Homo sapiens*). The ratio of the number of splice sites to the number of exons generally lay in the range of one to two (Figure 2A). This makes sense: if there were no splicing the number of splice sites would be zero, but the number of exons would be equal to the number of genes (i.e. the ratio would be zero); if there were a single intron, the number of splice sites would be two (one at the 5’ end of the intron and one at the 3’ end) and the number of exons is two. Thus the ratio would be one. If there are a large number of exons, then the number of introns would be nearly equal to the number of exons and so the number of splice sites is approximately two per exon (one at each intron end); in this case the ratio would be two. Thus, our splice site centric approach generalizes the exon-centric view. The advantage is that it allows us to analyze alternative splicing across whole transcriptomes, without relying on the annotation of exons.

**Figure 2.**
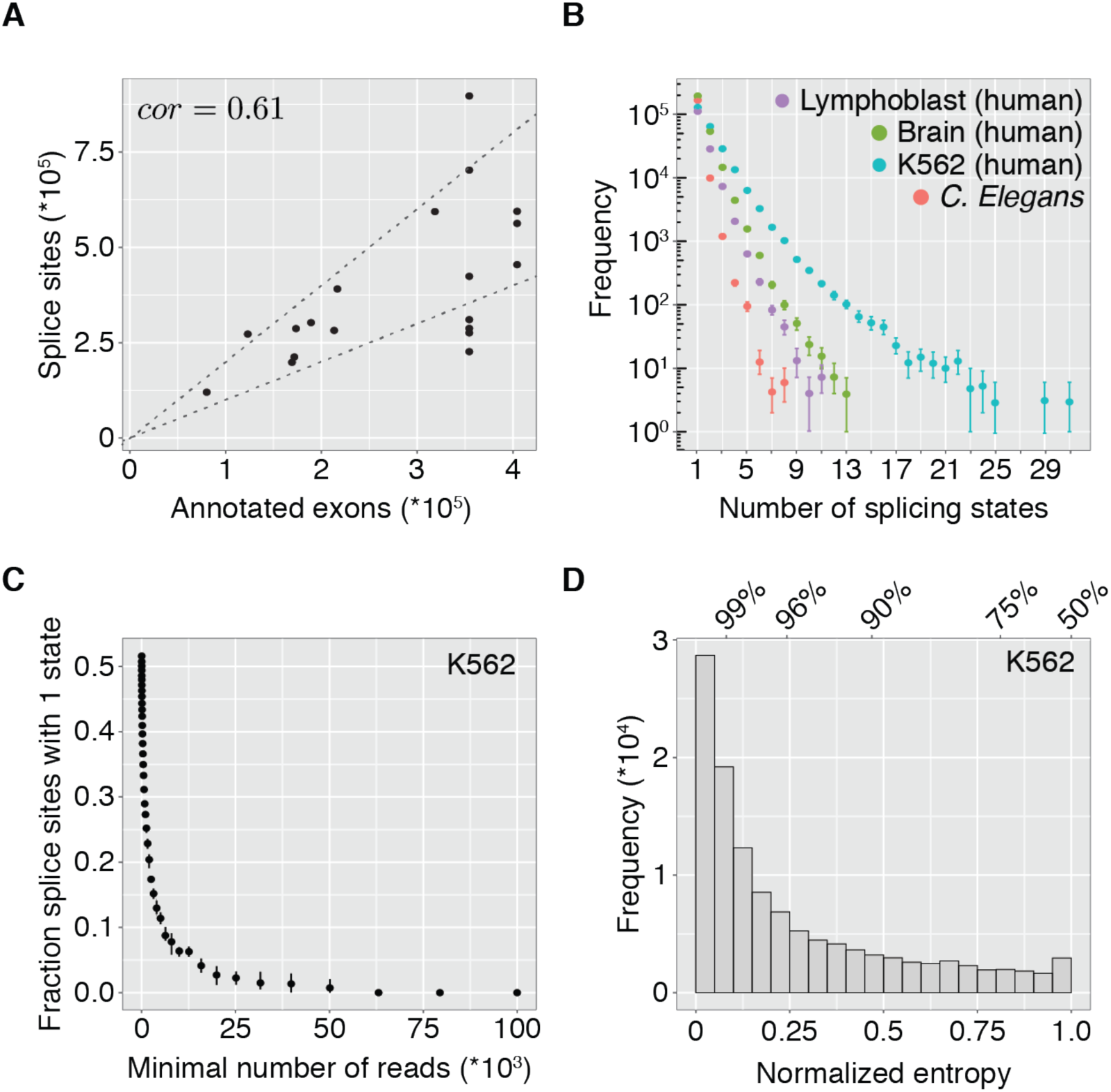
The majority of splice sites use only few choices of which one is used dominantly. **(A)** Number of splice sites identified by the splice-site centric approach versus number of exons annotated. Value pairs for each of 23 transcriptomes analyzed here are given. Dotted lines indicate linear functions with slope 1 and 2. Pearson correlation value between both sets is given. **(B)** Splice site count (Frequency, log10) versus number of splicing states for 4 transcriptomes with distinct expected complexity: *C. elegans* development (red), a human lymphoblast cell line (pathan, purple), the human cancer cell line K562 (blue) and human brain (green). See Supplementary Table 1 for sample details. Error bars represent 95% confidence intervals inferred by bootstrapping. **(C)** Fraction of splice sites with a single splicing state is determined for all splice sites with increasing minimal number of reads. Error bars represent 95% confidence intervals inferred by bootstrapping. **(D)** Normalized entropy histogram for the K562 transcriptome. The ratios in a 2-state system (i.e. minor and major splicing state) yielding the corresponding normalized entropy are plotted (top of panel), indicating non-linear behavior.

### The majority of sites use few splicing states

In each data set, we counted the number of splicing states (*N*_*i*_) associated with each splice site (i), where *i* runs from 1 up to the number of splice sites. From this, we determined the frequency histogram of the number of splicing states. Histograms were constructed for all 23 data sets; four histograms sets are shown in Figure 2B. For all 23 data sets, the most frequently observed number of splicing states is 1; the histogram of number of splicing state is monotonically decreasing; and the histogram can be approximated by an exponential function (figure supplement 1 & 2). The decay rate was fastest for *C. elegans*, 2.57, meaning that there is a 13-fold (e^2.57^) decrease in probability with each increasing number of states.The rapid decay indicates that there are relatively few highly alternatively spliced sites. The decay was slowest for the human K562 cell line, 0.7, meaning that there is a 2-fold (e^0.7^) decrease in probability with each increasing number of states. The slow decay indicates that there are relatively many highly alternatively spliced sites. The common characteristic of all the splicing state distributions is that they decrease monotonically as the number of splicing states increases. Thus the most common number of splicing states is one (*N*_*i*_ = 1) and those that are alternatively spliced generally have a small number of splicing states (the difference between the different data sets will be discussed later).

### There is no evidence for constitutive splicing

We found that the most common number of splicing states is one (i.e. *N*_*i*_ = 1). If constitutive splice sites are defined as having only one splicing state, then our analysis appears to support the conclusion that constitutive splicing dominates over alternative splicing at individual splice sites. However, a closer inspection of the data shows that this conclusion is likely to be wrong. We formed subsets of the data in which the number of reads exceeded a minimum number. For the full data set this number is 10, as described in the methods. For the K562 data set, there are splice sites with over 100,000 reads. When we plotted the fraction of splice sites with *N*_*i*_ = 1 as a function of the minimum number of reads, we found that the fraction decreased monotonically and approached zero. Furthermore, the frequency histograms constructed with data where the minimum number of reads is large (2500 in the case of K562 cells) were not monotonic but (figure supplement 2L). The important conclusion is that the fraction of splice sites with only one state (i.e. constitutive) depends on the number of reads and is small when the number of observations is large. Thus, we find no evidence for a separation between alternatively and constitutively spliced sites. The functional implications of this finding are discussed later.

### The majority of splice sites use one splicing state dominantly

Although the distribution of state counts allows us to characterize the availability of alternative splice site choices, it does not quantify how evenly these alternative states are used; for example, it does not distinguish between two-state systems that have probabilities 0.5/0.5 and 0.95/0.05. To quantify how information is expanded we use the information theoretic concept of Shannon entropy (Shannon, 1948):

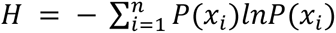

For example, two-state systems with probabilities 0.5/0.5 and 0.95/0.05 have entropies ~0.69 and ~0.20 respectively. The advantage of using entropy here is that it is a summary statistic capturing how evenly the available states are used.

As a first application for Shannon entropy is to address the question how evenly distributed splicing is among the alternative splicing states. Toward this end, we computed the normalized Shannon entropy (*H*/ln (*n*)). The normalized Shannon entropy is one if all splicing states are used at the same frequency; it is zero when a single state is used exclusively. We computed the distribution of normalized Shannon entropy values for the K562 transcriptome for all alternative splice sites in the transcriptome (Figure 2D and figure supplement 3). The average normalized entropy was 0.138, which for two isoforms corresponds to probabilities of 96.9% and 3.1% respectively. The average normalized entropy varied between transcriptomes, ranging from 0.028 in *D. rerio* (corresponding to probabilities of 99.6% and 0.4%) to 0.151 in *A. carolinensis* (corresponding to probabilities of 96.5% and 3.5%). Thus, most splice sites in all transcriptomes analyzed show low levels of normalized entropy, indicating that one splicing state, the major state, is predominantly used compared with all other states.

### Alternative splicing can be well approximated with a two-state model with a single alternative splicing probability

In a second application of Shannon entropy, we found that the second most abundant (minor) splicing state dominates over all other minor states. To show this, we approximated the full multinomial distribution for each splice site with a binomial distribution, considering only major and minor splicing states. We found that the Shannon entropy of the approximated system captured 85% of the information contained in the full system over the entire transcriptomes (Figure 3A and figure supplement 4). The data indicates a one-to-one correspondence between the information contained in the original data and its approximation (the log-log slope is 0.93). Thus most of the information is contained in the probabilities of the major and minor splicing states.

**Figure 3.**
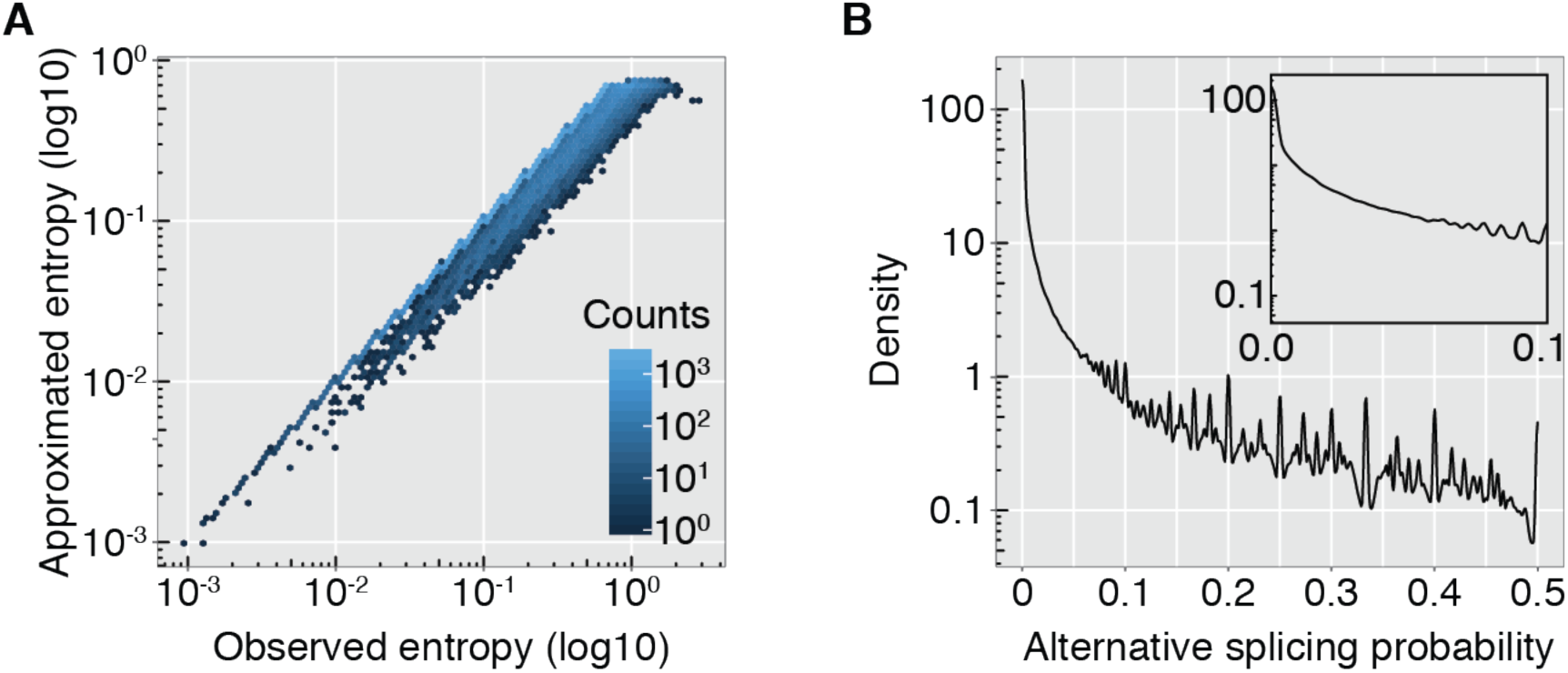
Alternative splicing can be well approximated by a two-state model with a single alternative splicing probability. **(A)** Entropy of the approximated two-state (binomial) model versus observed entropy for the K562 transcriptome. Value pairs are binned and counts per bin color-coded (log10). **(B)** Log-density plot of alternative splicing probability for the K562 transcriptome. Densities for low alternative splicing probabilities are highlighted (inset).

The advantage of the binomial approximation is that it allows us to represent each splice site by a single value, the probability of splicing to the minor splicing state. We call this value the alternative splicing probability, *p*_a_ (corresponding to *p*_2_ in Figure 1). Figure 3B shows the density histogram of alternative splicing probabilities for all splice sites in the K562 transcriptome. This distribution and analogous distributions derived from other transcriptomes (figure supplement 5) show rapidly decreasing density for large alternative splicing probabilities, reinforcing our finding that the major splicing state dominates over the minor one.

### The frequency of alternative splicing across transcriptomes follows a power law

To identify a frequency distribution function that describes the alternative splicing probability *p*_a_, we plotted a histogram of *p*_a_ on a log-log plot for three representative transcriptomes (Figure 4A). For all transcriptomes, there is a range of *p*_a_ over which the log-log plot is linear. This suggests that each *p*_a_ is distributed according to a power law:

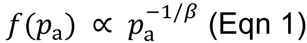

The parameter *β* has the following interpretation: a larger *β* implies *P*(*p*_a_) has a heavier tail and thus that there is more alternative splicing. We call *β* the extent of alternative splicing because the larger its value, the more extensive is alternative splicing.

**Figure 4.**
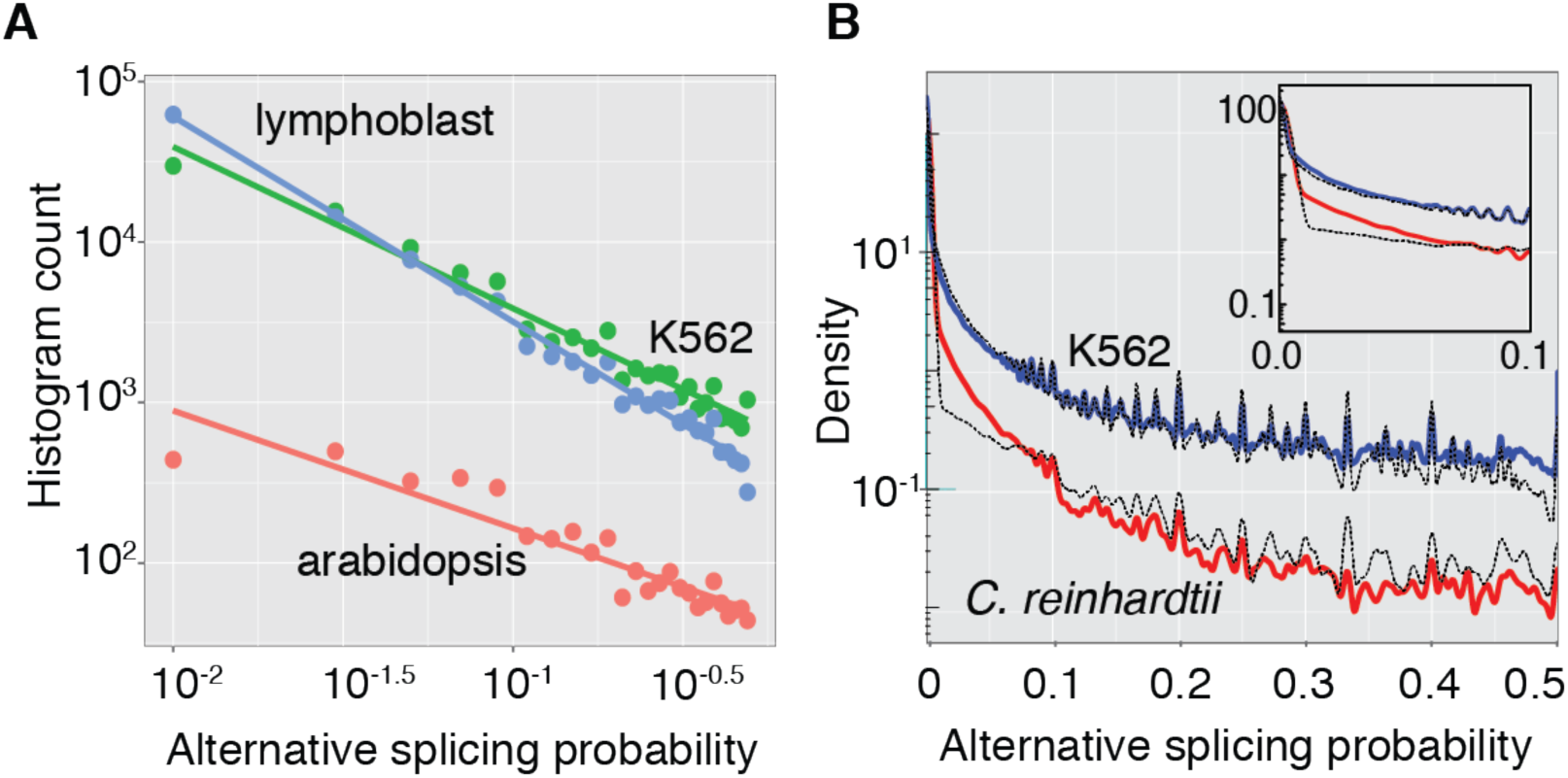
The probability of alternative splicing across transcriptomes is described by a power law model with a single free parameter, the extent. **(A)** Three representative histograms (*n* = 30 bins) of alternative splicing plotted on log-log axes reveals a linear regime; ordinary least squares fit to log-log data is also plotted. **(B)** Comparison of model and data for K562 (human) and *C. reinhardtii* transcriptomes (see figure supplement 8 for other transcriptomes). We plot the probability density function of the alternative splicing probability p_a_ for both the data (in blue and red, respectively) and the power law model (for both experiments, in black) on log-linear axes after performing a kernel density estimation. The model, which describes the probability of alternative splicing as a power law, reproduces many of the characteristics of the data for K562 cells and *C. reinhardtii*, two experiments of expected differing complexity.

To determine whether the power law provides a quantitative description of the measured alternative splicing densities, we took into account that each splice site has a finite number of reads (*n*), leading to visible peaks at certain ratios (Figure 3B). For example, if there are 10 total reads for a given splice site, then its associated *p*_a_ is necessarily 0.1, 0.2, 0.3, 0.4 or 0.5.

To take the discrete nature of the data into account, we generated a histogram of the model, which we then compared with the data. The histogram was generated as follows: for each splice site, we calculated an alternative splicing probability *p*_a_ from the power law (Eqn 1), see SI for the normalization), and then used the total number of read (*n*) to generate the number of minor reads given by the model from the binomial distribution *B*(*n*, *p*_a_). For each transcriptome, we estimated the exponent *β*, along with confidence intervals on this estimate, using the framework provided by Bayesian parameter estimation (see Materials and Methods and SI). Using this procedure, we found good agreement of the theoretical curves with the experimental data, including the visible peaks arising from the discrete sampling (Figure 4B).

The power law model provided a good description of *p*_a_ for all transcriptomes (figure supplement 6). The model recapitulated the salient aspects of the data, such as initial values (*P*(*p*_a_ = 0)), slopes and the discrete sampling effects discussed above. Furthermore, the power-law model explained more than 99% of the variation in the data, as measured by the coefficient of variation *R*^2^ (Table 1).

**Table 1.**
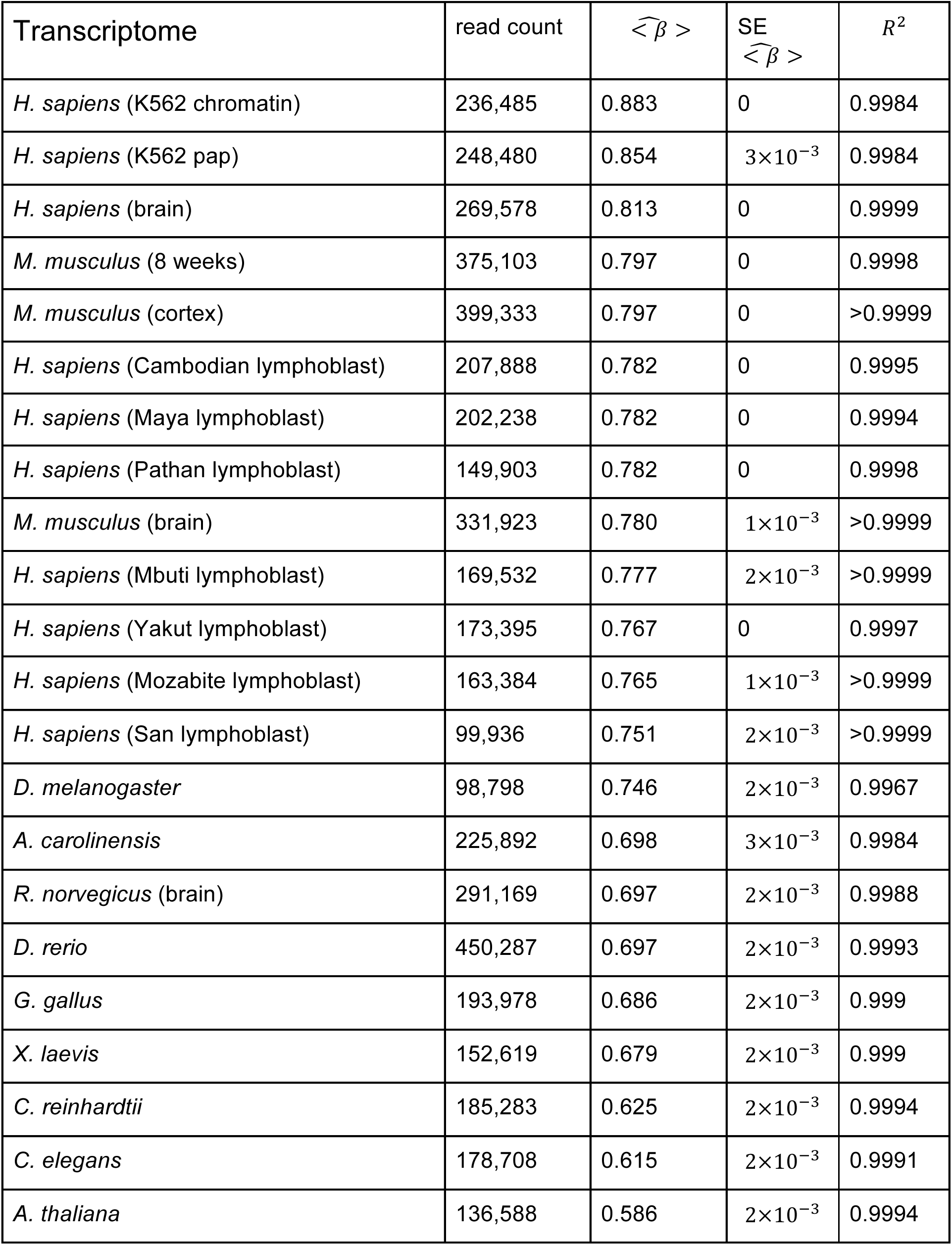
Transcriptome specific alternative splicing extent. As a function of transcriptome, we tabulate i) total number of splice sites, ii) predicted power law extent of alternative splicing, iii) standard error on the predicted power law extent (*n*=10) and iv) coefficient of determination. Transcriptomes “Cambodian”, “Maya”, “Pathan”, “Mbuti”, “Yakut”, “Mozabite” and “San” are all RNA-seq experiment on human samples from differing regions of the globe (Supplementary Table 1). For each transcriptome, the coefficient of determination was computed *n* = 20 times (due to the stochasticity of the process of simulating the model) and the largest SEM was 3.4×10^‒3^.

The extent of alternative splicing varied among transcriptomes. The minimum value was 0.58 for *A. thaliana* and the maximum 0.88 for K562 chromatin with high values corresponding to a slow decay, meaning more alternative splicing. The extents for human and mouse transciptomes clustered around 0.8. The extents for plants were considerably smaller (*C. reinhardtii* was ~ 0.63 and *A. thaliana* 0.58). The extents of *Drosophila* and lower vertebrates were intermediate (Table 1). Thus, our power law analysis identifies systematic differences in splicing between different cells and organisms.

### Many different features contribute to alternative splicing

Power laws often arise in nature when many components or features contribute to a process (Reed and Hughes, 2002). To test whether this applies to alternative splicing, we used machine learning techniques to correlate alternative splicing probability for each splice site with a large number of genetic and epigenetic features associated with that splice site. The 14 genetic features included the strength of splice site sequences, intron lengths and the distance between competing splice sites. For epigenetic features, we took advantage of the large amount of data on K562 cells from the ENCODE project, which measured chromatin context as well as the profiles of DNA and RNA binding proteins to provide 784 features (Figure 5A and supplementary Table 2). We further included splice site expression (the number of observations per splice sites) as the 785^th^ epigenetic feature.

**Figure 5.**
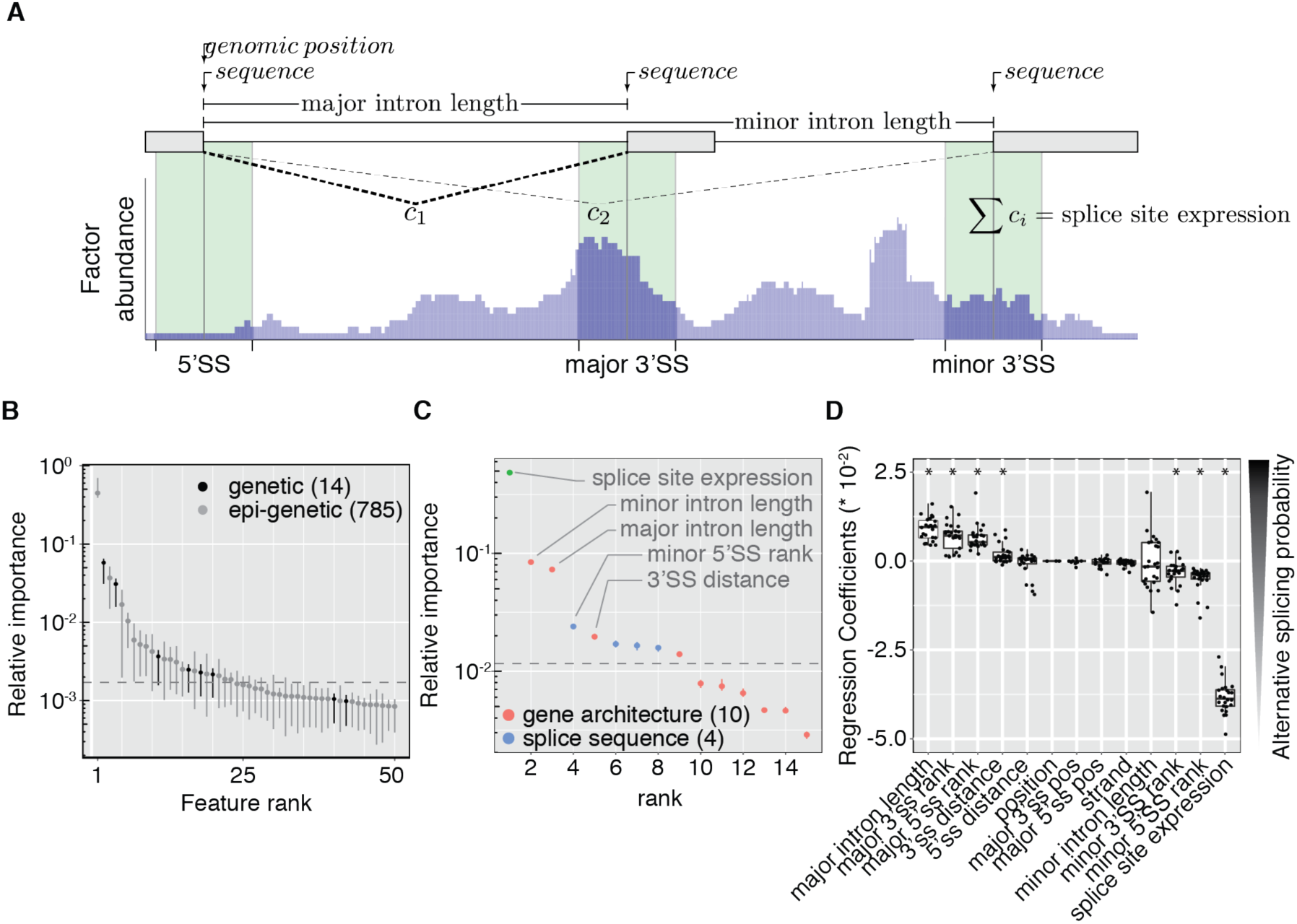
Many different features contribute to alternative splicing probabilities. **(A)** Schematic of feature extraction for an example splice site. The splice site (5’SS) as well as most abundant (major) and second most abundant (minor) splicing states are characterized by genetic (top part) and epigenetic features (bottom part). Genetic features include position, splice site strength and intron lengths. Epigenetic features include total number of reads per splice site (splice site expression) and features derived from genome-wide experiments performed in the ENCODE project. Factor abundance profiles (blue histogram) are intersected with windows up-and downstream of splice site of interest (green boxes) and mean abundance levels calculated. 14 genetic and 785 epigenetic features were determined for > 1.2 * 10⌃5 splice sites of the K562 transcriptome. **(B)** Ratio of observed and expected (by random choice) number of features in set of significant features (Enrichment, log10). Genetic and epigenetic features are enriched 14.9-fold and 0.75-fold respectively (p-values < 10⌃-5, left columns). Enrichment of functional subsets of epigenetic and genetic (right columns). Observed number of features within the groups of transcription factors (p-value < 0.05), gene architecture (p-value < 0.005) and splice sequences (p-value < 0.05) deviate significantly from values expected by random allocation. **(C)** Relative importance of features (Log10) to predict K562 p_as_ plotted versus rank. Feature importance was derive from random forest model containing only gene expression (green) and genetic features. Genetic features were divided into gene architecture (red) and splice sequence (blue) groups. The dotted line represents a 5% FDR cut-off determined by randomly generated features. **(D)** Regression coefficient are visualized for features with at least 3 data-points. Each data-point corresponds to a regression coefficient for one transcriptome and summary statistics are displayed by boxplots. Horizontal bars indicate median, boxes indicate interquartile range. Lasso regression was performed on all 23 transcriptomes and only algorithmically selected significant features displayed. Absolute value and sign of the regression coefficient indicate strength and direction of correlation with alternative splicing probability (right axis label).

We used a random forest non-linear regression model to predict alternative splicing probabilities (figure supplement 7A, *R*^2^ = 0.34) and identified 23 significantly enriched features (Figure 5B and supplementary Table 3). The most important features were splice site expression, major and minor intron length, Pol II occupancy, chromatin structure, histone modifications and splice site strength. Interestingly, 6 out of 14 genetic features contributed significantly. A random forest trained on a reduced feature set for K562 cells consisting of only genetic features and the splice site expression had similar predictive power compared to the full model which included the additional 784 epigenetic features (Figure 5C and figure supplement 7B, *R*^2^ = 0.32). Because of the dominant effect of splice site expression and the genetic features, we performed a simplified analysis on the other transcriptomes, which lacked detailed epigenetic data (figure supplement 8). For all transcriptomes, the dominant features were splice site expression, major and minor intron length, distance between competing splice sites and splice site strength.

Although random forests can determine the relative contribution of features, they cannot tell us whether a given feature contributes positively or negatively. To this end, we used a regularized linear regression model (lasso regression (Tibshirani, 1994), to predict alternative splicing probabilities for all transcriptomes. Feature importance is quantified by the absolute value of regression coefficients (Figure 5D) and agrees with feature importance quantified for random forest models: splice site expression is the feature with the highest importance, followed by intron lengths, splice site distance and strength. Some of the correlations accorded with biochemical expectations. For example a positive correlation between minor splice site strength and *p*_a_ and a negative correlation between major splice site strength and *p*_a_ accords with biological expectations (Sugnet et al., 2004). Other correlations, however, were unexpected. These included, a positive correlation with major intron length, indicating that distal splice sites are unfavorable for splicing of the major splicing state. Another unexpected finding was that splice site expression, the most important feature, is negatively correlated with alternative splicing probability indicating that highly expressed genes have little alternative splicing. These general findings were confirmed using a threshold-based analysis approach to classify splice sites into low (*p*_a_ < 0.1) and high (*p*_a_ > 0.1) alternative splicing groups (figure supplement 9). Thus three distinct analyses show that alternative splicing probabilities are influenced by a complex mixture of splice site characteristics. This is consistent with the notion that many components contribute to alternative splicing. Furthermore, our analysis offers molecular insights into the origin of the power law distribution.

### Alternative splicing extent contributes more to transcriptome expansion than exon count

Can the extent of alternative splicing explain the increase in transcriptome expansion? To quantify transcriptome expansion, we introduce the term diversity of a splice site to be *e*^*H*^, where *H* is the entropy (Eqn 1). The diversity equals the number of states required for the given entropy if all states are equally populated. Diversity corresponds to the ‘true diversity’, a term from population biology, which is used as an index of the diversity of an ecosystem (MacArthur, 1965). The extent of splicing correlates with the diversity averaged over all splice sites (Figure 6A). Thus, the broader the distribution of alternative splicing probabilities (i.e. the greater the extent of splicing), the higher the average diversity of splicing states.

**Figure 6.**
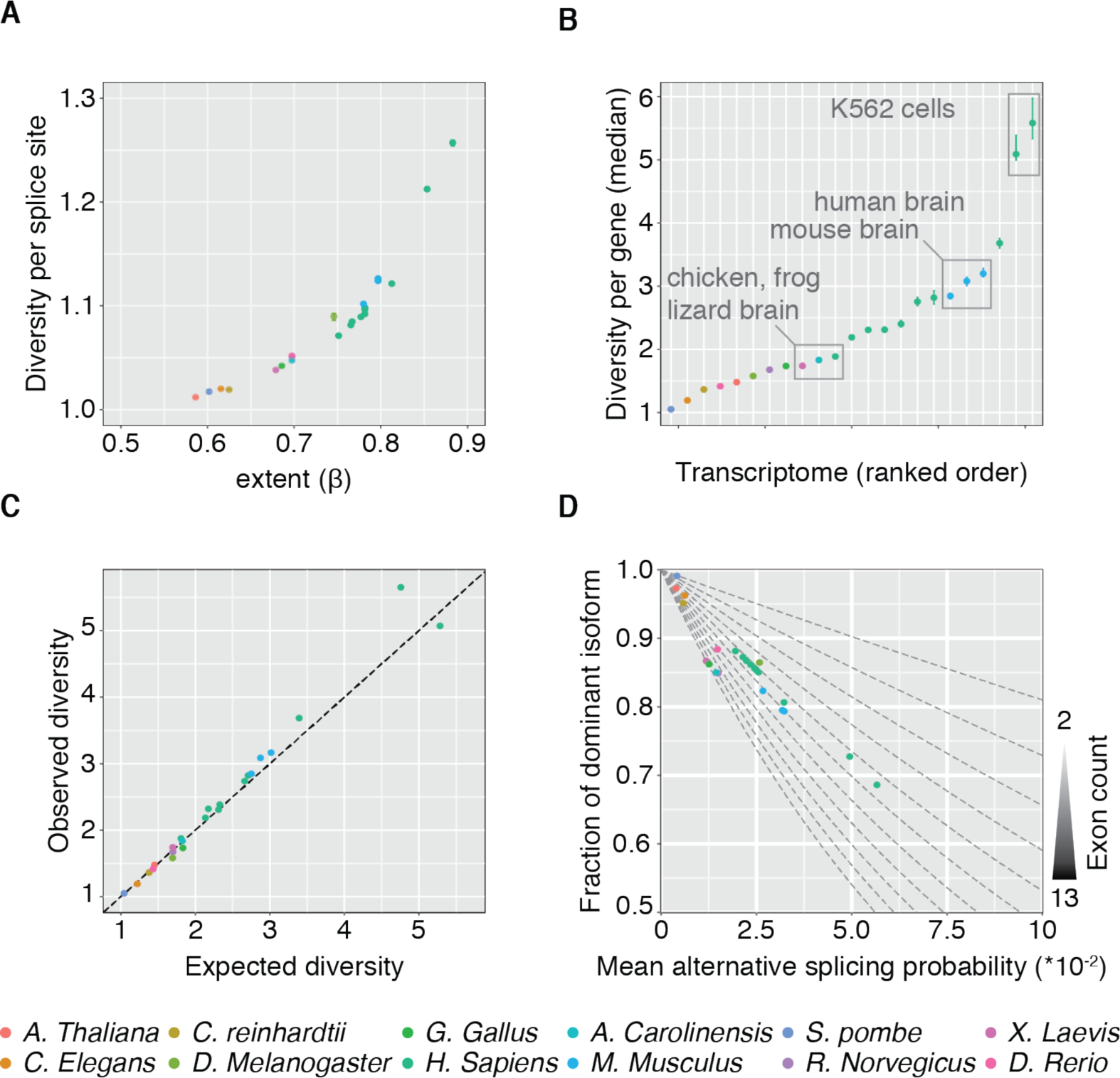
Alternative splicing probability combined with gene architecture determines transcriptome expansion. **(A)** Mean number of states effectively used per splice site plotted versus alternative splicing extent. Data-points represent a value pair for each transcriptome color-coded by species (legend at the bottom). **(B)** Median number of states effectively used per gene plotted in ranked order. Number of effectively used states was calculated from gene associated splice sites, assuming independency. Error bars represent 95% confidence intervals inferred by bootstrapping. Transcriptomes are color coding by species (legend at the bottom). **(C)** Median effectively used splicing states per gene (observed) plotted versus median effectively used splicing states per gene assuming random re-allocation of splice sites to genes. Linear function with slope 1 (dotted line) indicates a nearly 1 ‐to-1 correspondence. Each data-point represents one transcriptome. **(D)** Fraction of dominant isoform per gene expressed plotted versus mean alternative splicing probability per splice site. Available values for genes with S - 13 exons (label on right axis) are shown as dotted lines. Data points represent actual values for transcriptomes analyzed, derived from observed mean exon count and mean alternative splicing probabilities.

While the differences in the extent of alternative splicing (as measure by *β* or diversity) between transcriptomes are small (0.88S In K562 chromatin associated RNA to 0.586 in *A. thaliana*), these differences can lead to large differences in transcriptome expansion when there are multiple exons. If we assume independence of alternative splicing choices, then we can calculate diversities of the isoforms generated by each gene (i.e. over all its splice sites) as the sum of the diversities of the constituent splice sites. This is an upper limit of a gene’s true diversity and provides a quantitative measure of transcriptome expansion, based on information theoretic concepts, that takes into account the uneven distribution of isoform distributions.

Transcriptome expansion, as defined here, varies over a wide range for the 17 transcriptomes analyzed (Figure 6B). Transcriptome expansion ranged from 1.2 in *C. elegans* to ~1.75 in *G. gallus* brain to ~S.65 in *H. sapiens* brain. Interestingly, the cancer tissue culture cell line (K562) has by far the largest transcriptome expansion (~ 5.5) despite being only a single cell type. Note that transcriptome expansion is not dominated by a small number of genes that concentrate highly diverse splice sites: If alternative splicing probabilities were randomly allocated to genes an almost identical transcriptome expansion would be obtained (Figure 6C). Thus, combinatorial choice does indeed result in more pronounced transcriptome expansion.

Which of the two variables, alternative splicing extent and exon count contribute more to transcriptome expansion? Mean exon counts per transcript range from S.16 in *S. pombe* to 12.99 in *X. laevis* (figure supplement 10). Alternative splicing complexity is characterized by the extent of alternative splicing ranging from 0.59 for the least complex transcriptome (*A. thaliana*) to 0.88 for the most complex one (K562 chromatin associated RNA). The latter transcriptome represents the most complex one characterized here, achieving a median transcriptome expansion of 5.5 per gene. To achieve the same expansion with an intermediate (*β* = 0.70, *R. norvegicus*) or the minimal (*β* = 0.59, *A. thaliana*) observed alternative splicing extent would require the mean number of exons to increase from 7 to S5 or 7 to 140, respectively. While these increases in mean exon count are not observed, the observed range of power law exponents enables a wide range of transcriptome expansion factors. We conclude that transcriptome expansion is achieved mainly by varying alternative splicing extent.

How can expression of a functional dominant isoform be realized, given the alternative splicing complexity in combination with multi-exon genes? To address this point, we determined the fraction the major RNA isoform represents from all isoforms generated from the average gene. Figure 6D shows this fraction plotted against a range of mean *p*_a_ (0 to 0.1) for varying number of exons (2-1S). As expected, increasing the probability of alternative splicing decreases the fraction of the dominant isoform. This decrease is linear for genes with two exons (one splicing decision) and faster for increasing number of exons. Importantly, the observed mean exon counts and mean *p*_a_ implies that for all transcriptomes, the most abundant isoform has a relative abundance of > 70 % (Figure 6D, data points). For human transcriptomes, the median relative abundance of the most abundant isoform was ~85 %. Thus, while the majority of resources invested into the expression of the average gene goes to a major, functional isoform, ~15 % goes to minor isoforms. Non-zero alternative splicing probabilities for all splice sites (see above) result in a combinatorial expansion of the transcriptome.

## Discussion

We have developed a framework based on information theory that allows us to quantify how alternative splicing leads to transcriptome expansion. The framework is based on the concept of diversity (corresponding to “true diversity” in ecology (MacArthur, 1965), which takes into account both the number of possible RNA isoforms and how uniformly these isoforms are distributed. Using this measure, a splice site that has two splicing states of equal probability contributes more to transcriptome expansion than a splice site in which one state completely dominates over all other states. Taking splicing probabilities into account is essential because we find that, as the number of sequencing reads of a splice site increases, the chance that there is only one splice state (i.e. constitutive splicing) becomes very small. If we simply counted isoforms, the expansion would depend critically on the number of reads; instead, diversity provides a measure which is nearly independent of the density of transcriptome reads.

Provided that the splicing probabilities at each splice site in a gene are independent, we can calculate the expansion of the transcriptome as the average splice site diversity multiplied by the average number of splice sites per gene. If the probabilities are not independent, then our measure will overestimate expansion. Using this measure, we can estimate separately the contributions to transcriptome expansion of increased exon number and increased alternative splicing probability. To compare the relative contributions of splicing and exon number, consider K562 cells, which have the greatest transcriptome expansion (5.5) of all seventeen transcriptomes in our dataset. If K562 cells had the same splicing diversity as *A. thaliana* (the transcriptome with the lowest diversity), then to achieve a transcriptome expanse of 5.5 would require increasing the average number of exons from 7 to 140, which is an order of magnitude larger than the average exon number found in any species. This finding, namely that in the absence of increased alternative splicing very large increases in exon number would be needed, applies to other highly expanded transcriptomes in our database (Table 1). Thus, the increase in the diversity of splicing contributes more than the increase in the number of splice sites to the expansion of the transcriptome and highlights the importance of alternative splicing to transcriptome expansion.

Our information theoretic approach gives us a measure not just of the average effect of alternative splicing (through diversity), but also the distribution of alternative splicing probabilities through a concept that we call the extent of splicing. We show that a two-state model in which only major and minor states are considered (and the rare splicing states ignored) accounts for 85% of the additional information produced by alternative splicing. We find that for all the transcriptomes analyzed, the probability of the minor state (*p*_a_) is approximately distributed as a power law, meaning that, due to the heavy tail of a power-law distribution, higher *p*_a_ are more highly represented than in an exponential distribution (which falls off more quickly). We use the exponent of the power law to define the extent of splicing, which quantifies the breadth of the splicing distribution: the greater the extent, the greater the fraction of splice sites that have more highly spliced minor splicing states. The extent is a useful concept as it allows the splicing distribution of an entire transcriptome to be characterized by a single number, facilitating the comparison of the importance of splicing in transcriptomes of different cells and organisms. The extent is closely related to the diversity of splicing. Indeed, we find that the diversity specifies the extent (Figure 6A). The extent of splicing varies from a minimum of 0.586 in *A. thaliana* to a maximum of 0.883 in K562 Chromatin. Extent values clustered into evolutionary groups. For example, all human and mouse transcriptomes had high values, whereas plants and single-cell organisms had low values. Thus, there are systematic changes in extent that correlate with the “complexity” of the organism or cell type.

An unexpected finding was that there is no “constitutive” splicing (i.e. use of a single splicing state), at least in K562 cells, which had the highest read counts. This finding is based on the observation that if there are sufficient reads, almost all splice sites will be found to have one or more alternative states. Thus, the classification of exons in K562 into constitutively and alternatively spliced (Barbosa-Morais et al., 2012; Merkin et al., 2012) is not supported by our data. Rather, our K562 data support the concept that all splice sites are susceptible to alternative splicing, functional or otherwise. Whether this finding made on a cancer cell line generalizes to other, perhaps more well-regulated, transcriptomes will require deeper sequencing. We hypothesize that alternative splicing of splice sites with small *p*_a_ is a form of splicing “noise”. By noise, we mean alternative splicing which results in a transcript with no immediate functional role in the cell. Such splicing noise could result, for example, from occasional error in the splice site recognition by the spliceosome. The absence of peaks in the *p*_a_ distributions (Figure 3B and figure supplement 5) suggests that this noise is not constant over all splice sites. This indicates that some splice sites are more prone to mis-splicing than others. This is in contrast with the sequence independent noise in other gene expression processes, such as transcription and translation, which are characterized by a fixed error rate for all substrates. If we attribute the occurrences of minor splice site reads for splice sites with many reads (> 5 * 10^4^) solely to splicing error, then we estimate that the average error rate is 1 per 5000 splicing events. Note that both noise and functional alternative splicing contributes to this value and that it therefore is likely too large. Given that the average exon size is ~150 nt, the error rate corresponds to 1 per 7.5 * 10^5^ nt (or 1: 2.5 * 10^5^ amino acids). Even assuming that a mis-splicing event will destroy the function of the entire gene (which has several exons), this rate is small compared to transcription and translation error rates. Thus, we find that splicing, like other processes that mediate the flow of information from DNA to protein, is susceptible to noise, but that the errors introduced by splicing noise are smaller than those in other processes.

Noise in alternative splicing noise may confer evolutionary advantages to an organism. A low rate of noise creates many minor alternative isoforms. A role of these isoforms may be to extend the potential sequence repertoire evolution can work on. A very rare isoform is not expected to have a great influence on protein function. Therefore, it can be acted on by evolution without affecting the original function of the gene, allowing the cell to improve fitness without gene duplication (Boue et al., 2003; Gilbert, 1978). Importantly, a biological system has to strike a balance between expressing few functional major isoforms and many non-functional minor isoforms. If only functional isoforms are expressed, no exploration of the alternative splicing landscape is possible. If too many minor non-functional isoforms are expressed, there may be not enough transcripts of the functional isoform. Here we show that the power law distributed alternative splicing probability over multi-exon genes results in the expression of one dominant major isoform and many minor isoforms. Thus, this enables robust expression of functional transcript and exploration of the combinatorial alternative splicing landscape in parallel.

## Materials and methods

### Splice site centric quantification of alternative splicing

RNA-Seq data (see Supplementary Table 1) was mapped to the respective genomes using Tophat2 (Kim et al., 2013). The splice site centric alternative splicing quantification was performed by custom software (https://github.com/carrillo). In short, reads spanning distal genomic locations are extracted. Splice site locations are inferred and reads allocated to these sites. From these allocated reads, each splice site is characterized by chromosomal position, chromosomal position of splicing states and number of reads (observations) for each splicing state. For transcriptomes with multiple RNA-Seq data-sets (see Supplementary Table 1) splice site characterization of each RNA-Seq sample are combined into one. To reduce statistical fluctuations and thereby increase confidence in our characterization of alternative splicing, we only analyzed splice sites with ten or more reads. The sole exception for this approach is the analysis performed in Figure 2A. The median read number ranged from 31 to 746 in the different data sets.

### Information theoretical approach to alternative splicing

Each splice site is characterized by a multinomial probability distribution, inferred by the frequency of splicing state observations. Information content of each splice site is measured by calculating the Shannon entropy from this probability distribution:

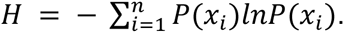

Shannon entropy values are not directly comparable for splice sites with distinct number of states. To achieve comparability, normalized entropy values are calculated:

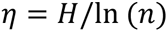

To transform the measure of entropy back to a measure of state-count we use the concept of true diversity:

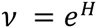

This measure generalizes the range of a probability distribution by weighing events by their probability. If all events are equally probable the extent equals the range. If *n* events are dominant it asymptotes to *n*. To deduce the information content of genes, we used available transcriptome information to allocate splice sites to genes. The information content of genes is calculated as the joint entropy of its associated splice sites. Importantly, this calculation assumes independence of splice sites and therefore provides an upper entropy bound. Effective state counts of genes are calculated from this joint entropy.

### Probabilistic Modeling of Alternative Splicing

We wish to model *p*_a_, the probability of alternative splicing. For any given splice site *S*_*i*_, *p*_a_(*S*_*i*_) = *n*_*i*_/*k*_*i*_, where *n*_*i*_ is the total number of reads of *S*_*i*_ and *k*_*i*_ is the number of observations supporting the minority state. We used a model in which the binomial probability is distributed according to a power law (see SI for other approaches). We define the model as follows: assuming a given extent *β* and exponent 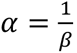 for each (*n*_*i*_, *k*_*i*_)-pair we first generate a *p* from 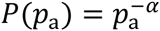; next, we use the generated *p* to generate a number of minority reads *k*_*i*_ from the binomial distribution with binomial probability *p*, B(n, p). Thus, given the experimental data of (*n*_*i*_, *k*_*i*_)-pairs, for a given exponent, we are able to simulate the model distribution.

We need to estimate the parameter *β* for each transcriptome in light of the available data and do so using Bayesian inference (see SI). Due to the size of the data sets in question, the posterior is dominated by the likelihood function *P*(*D*|*α*)and independent of the prior distribution. Thus the results reported are independent of the prior chosen (see SI) and rely on only the data and no prior assumptions. Performing a parameter sweep by varying *β* allows us to calculate the posterior mode. We eventually performed the parameter sweep for *α* on the interval [1.01,1.99] divided into 41 points (we did this as values *α* > 2 consistently gave poor fits for representative data sets). We retrieved estimates < *β̂* >, along with confidence intervals on these estimates (Table 1).

### Machine learning of features correlating with alternative splicing

Genetic features (Supplementary Table 2) were extracted from publicly available genome sequence and annotation files by custom code (https://github.com/carrillo). Epigenetic features were extracted from publicly available experiments conducted in the ENCODE project (Consortium, 2012). Raw data was mapped to the respective transcriptome using Tophat2 (Kim et al., 2013) and genome-wide chromosome abundance profiles were generated by Samtools (Li et al., 2009) and intersected with regions of interest using custom software (https://github.com/carrillo). For all machine learning models employed here the following pre-preprocessing steps were used: splice sites with *p*_*a*_ = 0 were removed, *p*_*a*_ were logit-transformed (Lasso regression only), categorical features were one-hot encoded, missing values were imputed by the feature median value, features were Box-Cox transformed to approximate normality (Lasso regression only) (Box, 1964), centered (μ=0) and scaled (sd=1). Model parameter selection was performed on 80% of the data. Selection was performed by grid-search using 5-fold cross-validation using mean squared error (MSE) as performance measure. Random forest models were implemented in Python using the scikit-learn framework (Pedregosa et al., 2011). Random forests contained 512 trees and maximal tree depth and minimal samples per leaf were optimized by cross-validation. Lasso regression was implemented in R using caret and glmnet packages (Friedman et al., 2010; Kuhn, 2008) and optimal regularization parameter determined. Model performance was determined on the 20% hold-out set using the optimized model parameters.

## Author contribution

F.C.O., H.B-A. and J.H. designed the study and wrote the paper. F.C.O. designed and implemented the splice-site centric quantification. F.C.O. selected, collected and analyzed the transcriptome data. H.B-A. designed and implemented the probabilistic modeling. F.C.O. designed and implemented the machine learning and integrated splice-centric quantification with genome architecture.

## Acknowledgments

We thank Karla M. Neugebauer, David Stanek, Julien Berro, Stephan Preibisch and Mark Gerstein and all members of the Howard group for discussions and critical comments on the manuscript. This work was supported by funding from MPIPKS Dresden, MPI-CBG Dresden, Yale University.

## Glossary

Transcriptome: Set of all RNA molecules in a sample (e.g. cell, tissue, organism).
Transcriptome expansion: Increase of coding expansion of the genome.
Gene annotation: Meta information added to the raw DNA sequence, such as exon-intron structure.
Gene architecture: Exon-intron structure of genes.
RNA Splicing: RNA maturation event leading to removal of introns and joining of exons.
Intron: Sequence removed by splicing, often non-coding for proteins.
Exon: Sequence retained by splicing, often coding.
Splice site: Exon-intron (5’ splice site) or intron-exon boundary (3’ splice site).
Constitutive splicing: The process that results in the joining of two splice-sites in all observed situation.
Alternative splicing: The process that results that one splice site can be joined to distinct partner splice sites.
RNA-seq experiment: Qualitative and quantitative profile of transcriptome by deep sequencing.
Extent: A parameter used to characterize the amount of alternative splicing in any given transcriptome; technically, the extent 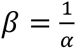, where *α* is the exponent in the power law distribution that describes the amount of alternative splicing in the transcriptome.
Splice site expression: Number of RNA-Seq observations per splice site.
Shannon Entropy: Metric of the expected information content.
True Diversity: An ecological concept which measures both the number of distinct species (richness) and how uniformly they are distributed in a sample (evenness).
Machine Learning: Computational algorithms which learn rules (model) to predict an output from an input.
Random Forest: A non-linear machine learning model based on an ensemble of decision trees with random feature subset selection at each decision node.
Lasso Regression: Linear regression regularized by absolute value of the sum of all regression coefficients (L1 norm).
Bootstrapping: Resampling technique to infer the confidence in a population measurement.
Probability density function (pdf): A function of a random variable X that describes the relative frequency for X to take each of its specific values.
Kernel density estimation: A method of estimating the probability distribution function based on a finite sample of data.
Bayesian Inference: A method of statistical reference in which prior knowledge is recursively updated utilizing new data using Bayes’ Theorem in order to make statements about probablistic hypotheses.
Prior distribution: The distribution (pdf) that mathematically formalizes one’s belief about the state of the system before taking empirical evidence (data) into account (note that the distribution can be a mathematical formalization of being in a state of ignorance).
Posterior distribution: The distribution that describes the probability of the random variable in question after the evidence/data is taken into account.

